# Why do plants make opioids? Testing the herbivore defense hypothesis for psychoactive alkaloids in kratom

**DOI:** 10.64898/2026.07.15.738691

**Authors:** Brent D. McNutt, Melissa R.L. Whitaker

## Abstract

A growing body of literature proposes that psychoactive plant compounds evolved as defenses against herbivorous insects, with their neurological effects in humans an evolutionary accident of conserved receptor architecture. This hypothesis – which we call the herbivore defense hypothesis – is widely invoked but rarely tested empirically. We report a controlled dietary bioassay in which *Spodoptera frugiperda* larvae were fed artificial diet incorporating lyophilized kratom (*Mitragyna speciosa*) leaf powder or purified mitragynine, the primary psychoactive compound in kratom leaf. Kratom leaf caused dose-dependent larval mortality, suppressed growth rates, and reduced pupation rates across all tested concentrations, with near-complete mortality at the highest dose. Purified mitragynine produced modest but significant growth suppression relative to the control, but had no significant effect on survival and was substantially outperformed by whole leaf powder at the same mitragynine-equivalent concentration. Because insects lack the *µ*-opioid receptors through which mitragynine exerts its psychoactive effects in mammals, its insecticidal activity cannot be mediated by the same receptor interaction responsible for its human pharmacology. We use these findings to critically evaluate the herbivore defense hypothesis, and find that its current framing is insufficient to answer one of chemical ecology’s most compelling questions: why do plants make compounds that alter the human mind?

## Introduction

Plants produce a remarkable diversity of secondary metabolites whose ecological functions have been debated since Fraenkel’s foundational observation in 1959 that herbivory shapes plant chemodiversity (1). Among the most pharmacologically potent of these are alkaloids – nitrogen-containing compounds that disrupt animal physiology with extraordinary specificity. Many plant alkaloids are psychoactive and used by humans as stimulants such as nicotine, caffeine, and cocaine; analgesics such as morphine, codeine, and mitragynine; and hallucinogens such as mescaline and dimethyltryptamine. A longstanding hypothesis proposes that plant alkaloids evolved as chemical defenses against herbivorous insects, the primary consumers of plant biomass and the principal selective agent on plant secondary chemistry (1; 2; 3). Under this framework, the neurological properties of plant alkaloids in mammals (including their diverse psychoactive effects in humans) are not the selected targets of these compounds but rather an evolutionary spandrel resulting from conserved receptor architecture (4; 5; 6). This hypothesis is conceptually attractive but rarely tested directly. Most arguments for a herbivore defense origin of psychoactive plant compounds reason backwards from pharmacological activity to presumed ecological function, without demonstrating that the compound actually affects herbivores at ecologically realistic doses.

Kratom (*Mitragyna speciosa* Korth., Rubiaceae) represents an unusually tractable system for testing the herbivore defense hypothesis. Native to Southeast Asia, kratom has attracted intense pharmacological interest due to the psychoactive and analgesic properties of its primary alkaloid, mitragynine, which acts as a partial agonist at *µ*-opioid receptors (MOR) in mammals (7). Despite a recent profusion of pharmacological and regulatory attention, the ecology of *M. speciosa* is almost entirely uncharacterized. Kratom contains mitragynine at 0.47% dry weight in juvenile leaves up to 1.6% in mature leaves (8), but the ecological function(s) of kratom secondary compounds has received almost no empirical investigation. A recent study by Leksungnoen et al. (9) documented eighteen potential insect herbivores in a kratom plantation in Thailand and found that insect-damaged leaves accumulate significantly higher mitragynine concentrations than undamaged leaves. This pattern is certainly consistent with inducible chemical defense and suggests that kratom alkaloid production may respond to herbivore pressure, but whether these alkaloids actually harm the insects that consume them has not been tested.

Here we report the results of a controlled dietary bioassay in which *Spodoptera frugiperda* (J.E. Smith; Lepidoptera: Noctuidae) larvae were reared on artificial diet incorporating kratom leaf powder or purified mitragynine. *Spodoptera frugiperda* and its congeners are highly polyphagous agricultural pests capable of feeding across hundreds of plant species (10; 11), and are among the most toxin-tolerant lepidopteran larvae known (12; 13), thus are widely used for testing insecticidal activity (14). The primary objective of this study was to evaluate the herbivore defense hypothesis by addressing a fundamental question in neuro-chemical ecology: do psychoactive compounds also function as anti-herbivore defenses?

## Materials and methods

Mature *M. speciosa* leaves were collected from a chemotyped and genotyped cultivar “Ms cv 2” (8) in cultivation at the University of Florida. After collection, leaves were lyophilized overnight, ground to a fine powder in a mortar and pestle under liquid nitrogen, then stored at -20^*°*^C until use.

*Spodoptera frugiperda* were hatched from a single egg mass (Benzon Research) and reared individually in thirty-two well plastic trays (Frontier Agricultural Sciences #9074) with breathable mylar lids in a Percival Scientific incubator at 27°C, 12:12 L:D cycle, and ambient humidity. Larvae were fed artificial diet (Frontier Agricultural Sciences, #F9772) incorporating pure mitragynine (Cayman Chemical #11151), lyophilized kratom leaf powder at three concentrations, or control diets. Whole leaf diets were prepared assuming approximately 1% mitragynine content by dry weight (8) and designated as ‘low’ (24.5 mg/g diet), ‘mid’ (49.1 mg/g diet), and ‘high’ (196.3 mg/g diet). These treatments were selected to include the range of naturally occurring concentrations of mitragynine in kratom leaves of different developmental stages (Table 1 (8; 15)). Pure mitragynine was dissolved in 1 mL of 70% ethanol and incorporated into prepared diet at 0.491 mg/g, a dose calculated to match the mitragynine content of the mid leaf diet rather than the high leaf diet due to the prohibitive cost of purified mitragynine. This dose nonetheless provides a direct internal comparison between whole leaf powder and its primary alkaloid at equivalent concentrations, and a solvent carrier control containing the equivalent volume of 70% ethanol was included alongside an untreated clean diet control to distinguish alkaloid effects from any solvent effects.

**Table 1.** Dietary incorporation rates and estimated mitragynine doses relative to naturally occurring concentrations across leaf developmental stages. ^*a*^Juvenile and mature leaf mitragynine concentrations estimated as 1.38 and 4.23 mg/g fresh leaf respectively, based on reported dry weight alkaloid content of 0.47% (juvenile) and 1.44% (mature) across leaf developmental stages (8) and a fresh-to-dry weight ratio of 3.4:1 (15)

| Treatment | Incorporation rate<br>(mg/g diet) | Mitragynine delivered<br>(mg/g diet) | % of juvenile leaf<br>concentration<sup>a</sup> | % of mature leaf<br>concentration<sup>a</sup> |
| --- | --- | --- | --- | --- |
| Low Leaf | 24.5 | 0.245 | 17.8% | 5.8% |
| Mid Leaf | 49.1 | 0.491 | 35.6% | 11.6% |
| High Leaf | 196.3 | 1.963 | 142% | 46.4% |
| Mitragynine | 0.491 | 0.491 | 35.6% | 11.6% |

Each larva was assigned to a single treatment, with thirty-two individuals per treatment group (n = 192 total). Larvae were fed *ad libitum*, given a ∼ 1cm^3^ cube of their assigned treatment that was replaced as needed. Daily checks recorded mortality and developmental milestones. Prior to pupation, larvae were transferred to 2oz lidded plastic deli cups. Pupal weight was assessed twenty-four hours after initial documentation of pupation. Adult eclosions were recorded at daily checks. The bioassay was considered complete when the last individual either eclosed or was confirmed dead. Pupae that remained in the pupal stage more than ten days beyond the last eclosion and showed no movement or response to stimuli were classified as pupal mortalities.

Survival and development data were analyzed using time-to-event methods, with time measured in days from hatch until death, pupation, eclosion, or the end of the observation period. Kaplan–Meier curves were estimated for each treatment group to visualize survival trajectories. Because the proportional hazards assumption was violated (Schoenfeld residuals, global *p* = 0.024), a log-logistic accelerated failure time (AFT) model was fitted as the primary parametric analysis, selected over Weibull, log-normal, and exponential alternatives by AIC. Treatment group was the sole predictor with the untreated control as the reference, so all time ratios reflect survival time relative to the untreated control. The mitragynine vs. solvent carrier control contrast, which isolates the alkaloid effect from the solvent vehicle, was addressed in planned pairwise log-rank tests rather than in the AFT model. A Cox proportional hazards model was fitted secondarily for descriptive purposes; hazard ratios are reported alongside their confidence intervals but are not interpreted as primary results given the violated assumption. To test for a monotonic dose-response relationship across ordered leaf concentrations, a log-rank trend test was implemented as a one-degree-of-freedom score test from a Cox model with a numeric dose score. Pairwise differences in larval survival were evaluated using five pre-specified log-rank tests with Holm correction. The same five comparisons were applied consistently across all endpoints (larval survival, pupal weight, and larval growth rate): low, mid, and high leaf vs. untreated control; mitragynine vs. its solvent carrier control; and mid leaf vs. mitragynine. Pupation and eclosion rates were compared across treatment groups using Gray’s test for cumulative incidence functions, which accounts for the competing risk of larval death before each milestone. For each individual that successfully pupated, larval growth rate was calculated as pupal weight divided by days to pupation to account for the confounding effect of prolonged larval duration on final body mass. Larval growth rate was compared across treatment groups using a Kruskal–Wallis test followed by the five pre-specified pairwise Wilcoxon rank-sum tests with Holm correction. All analyses were conducted in R version 4.5.1 (16) using the *survival* (17), *survminer* (18), *cmprsk* (19), *broom* (20), and *gtsummary* (21) packages.

## Results

Larvae fed on artificial diets spiked with whole kratom leaves or purified mitragynine differed substantially in their growth, survival, and development rates. A strong dose-dependent effect of leaf concentrations was observed on larval mortality: larvae from the high leaf treatment experienced an LT_50_ of four days (Fig 1), whereas all other treatment groups experienced *>* 50% survival for the duration of the experimental period. Larval survival differed markedly across treatment groups in a dose-dependent manner (log-logistic AFT model: *χ*^2^(5) = 81.47, *p* < 0.0001; *n* = 192). A log-rank trend test across ordered leaf concentrations confirmed a significant monotonic increase in larval mortality with increasing leaf dose (*χ*^2^(1) = 65.51, *p* < 0.0001). The log-logistic AFT model estimated that larvae in leaf-treated groups experienced substantially accelerated time to death relative to the untreated control: time ratios were 0.18 (95% CI: 0.03–0.96, *p* = 0.045), 0.12 (95% CI: 0.02–0.67, *p* = 0.016), and 0.02 (95% CI: 0.00–0.11, *p* < 0.0001) for low, mid, and high leaf, respectively, indicating that larvae in these groups died in 18%, 12%, and 2% of the time that control larvae would have survived. Planned pairwise log-rank tests confirmed that all three leaf concentrations differed significantly from the control group after Holm correction (low leaf: *χ*^2^ = 6.19, *p* = 0.039; mid leaf: *χ*^2^ = 12.55, *p* = 0.002; high leaf: *χ*^2^ = 53.43, *p* < 0.0001). The high leaf group experienced near-complete larval mortality, with 31 of 32 larvae dying before pupation; results for this group should be interpreted with caution given the effective sample size of one survivor. The solvent carrier control did not differ significantly from control in either the AFT model (TR = 0.34, 95% CI: 0.06–2.00, *p* = 0.234) or planned log-rank test (*χ*^2^ = 0.56, *p* = 0.453), indicating that the solvent vehicle alone did not substantially elevate larval mortality. Mitragynine likewise did not differ significantly from its solvent carrier control (*χ*^2^ = 0.56, *p* = 0.453), and the mid leaf vs. mitragynine log-rank comparison was not significant after Holm correction (*χ*^2^ = 3.56, *p* = 0.119), though this comparison was better powered in the growth rate analysis (see below).

**Fig 1.**
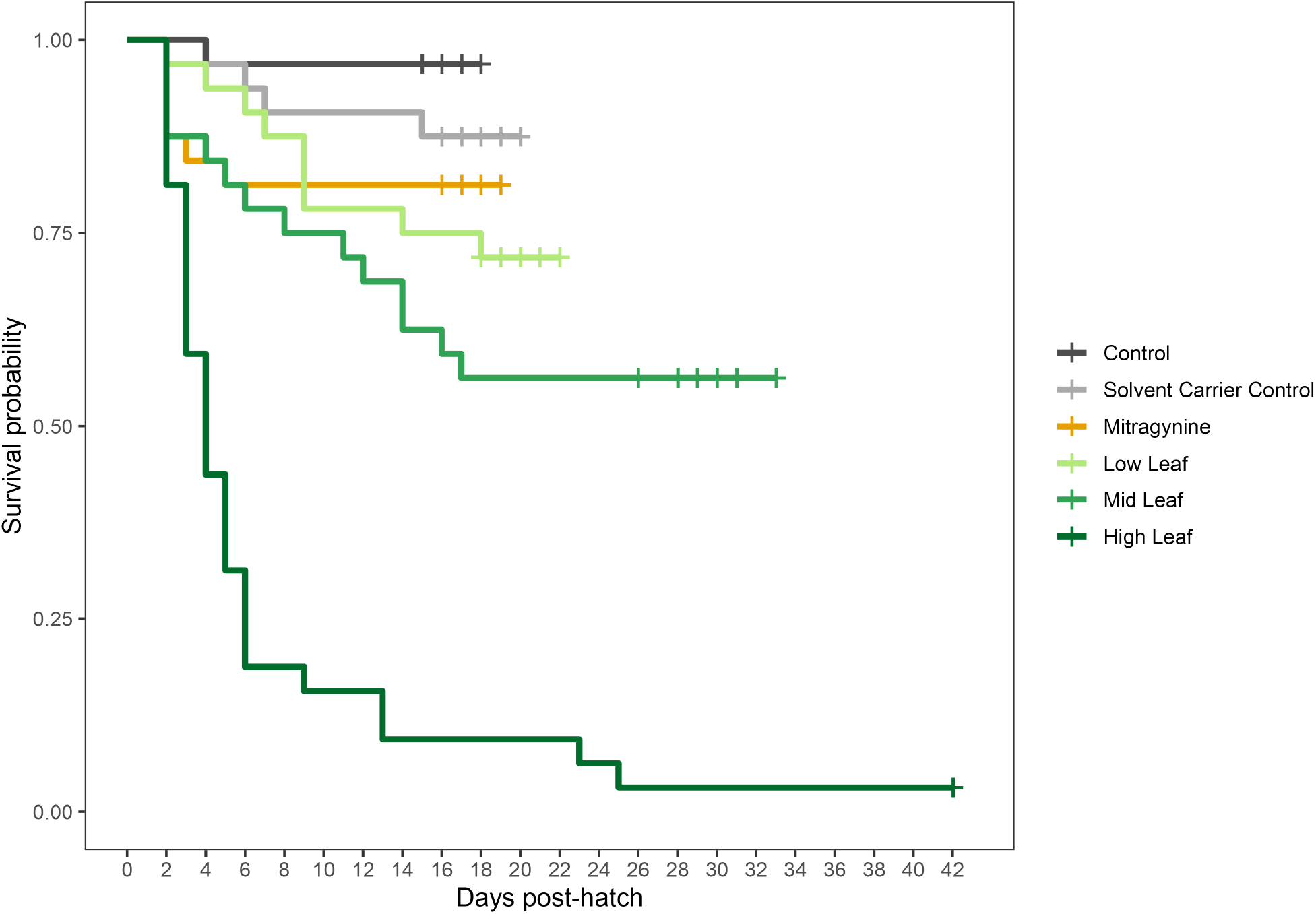
Larval mortality. Kaplan–Meier survival curves for *Spodoptera frugiperda* larvae across six treatment groups. Vertical tick marks correspond to censorship events (e.g. pupation); survival lines end when the last individual from each group died or pupated.

Time to pupation followed a similar dose-dependent pattern across leaf treatments (Fig 2). Among the control group, 96.9% of larvae pupated, followed by the solvent carrier control (87.5%), mitragynine (81.2%), low leaf (71.9%), mid leaf (56.2%), and high leaf (3.1%) groups. A Gray’s test confirmed that the distributions of time-to-pupation differed significantly across treatment groups (stat = 600.2, *p* < 0.0001), reflecting both delays in pupation timing and outright mortality before pupating. Cumulative incidence curves showed that control larvae reached peak pupation probability earliest (approximately days 14–18 post-hatch), with leaf-treated groups lagging increasingly behind. Only a single larva pupated from the high leaf treatment, at 42 days.

**Fig 2.**
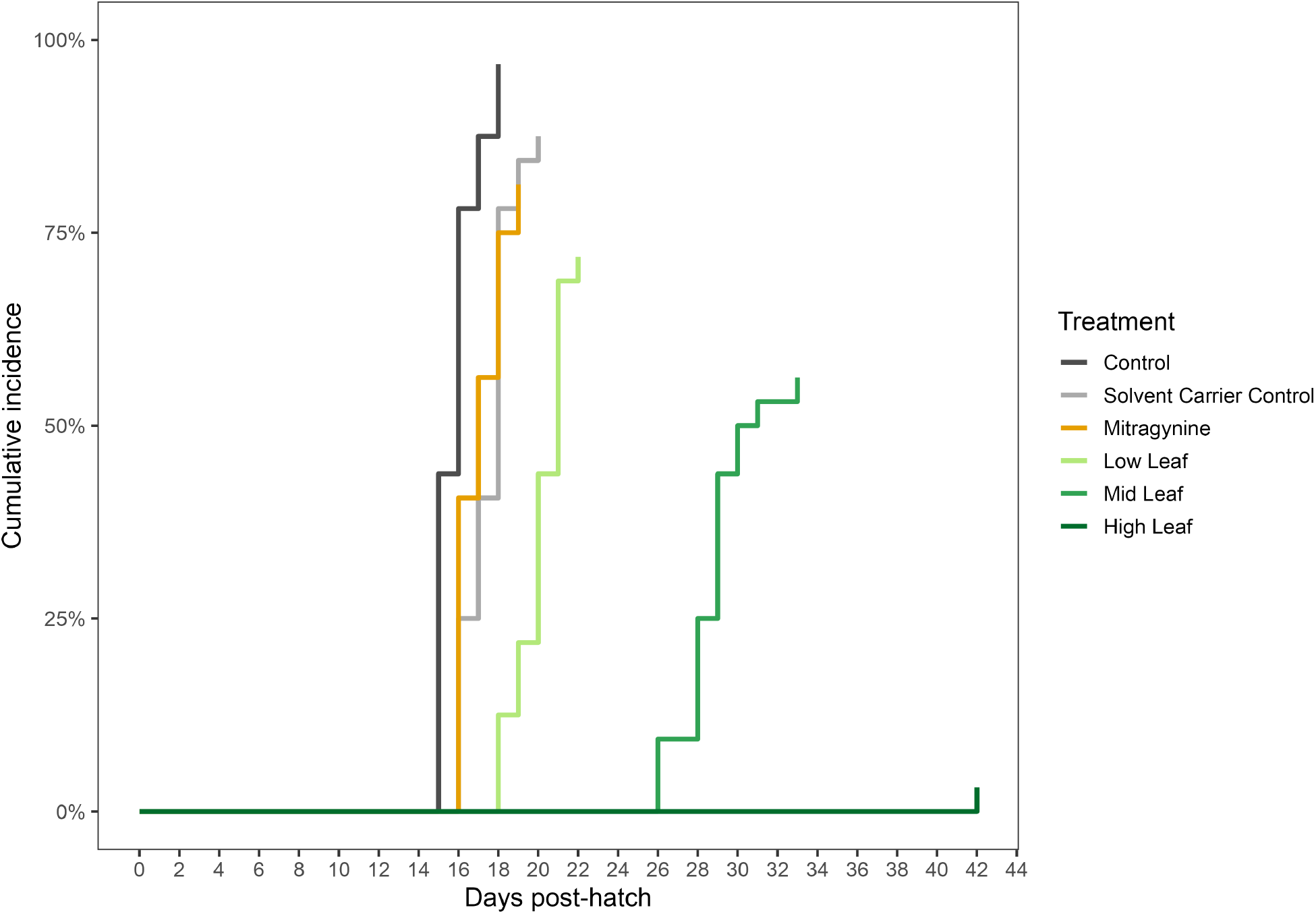
Time to pupation. Cumulative incidence of pupation across treatment groups, accounting for larval death as a competing risk. Note the single individual from the high leaf treatment that pupated at day 42.

**Fig 3.**
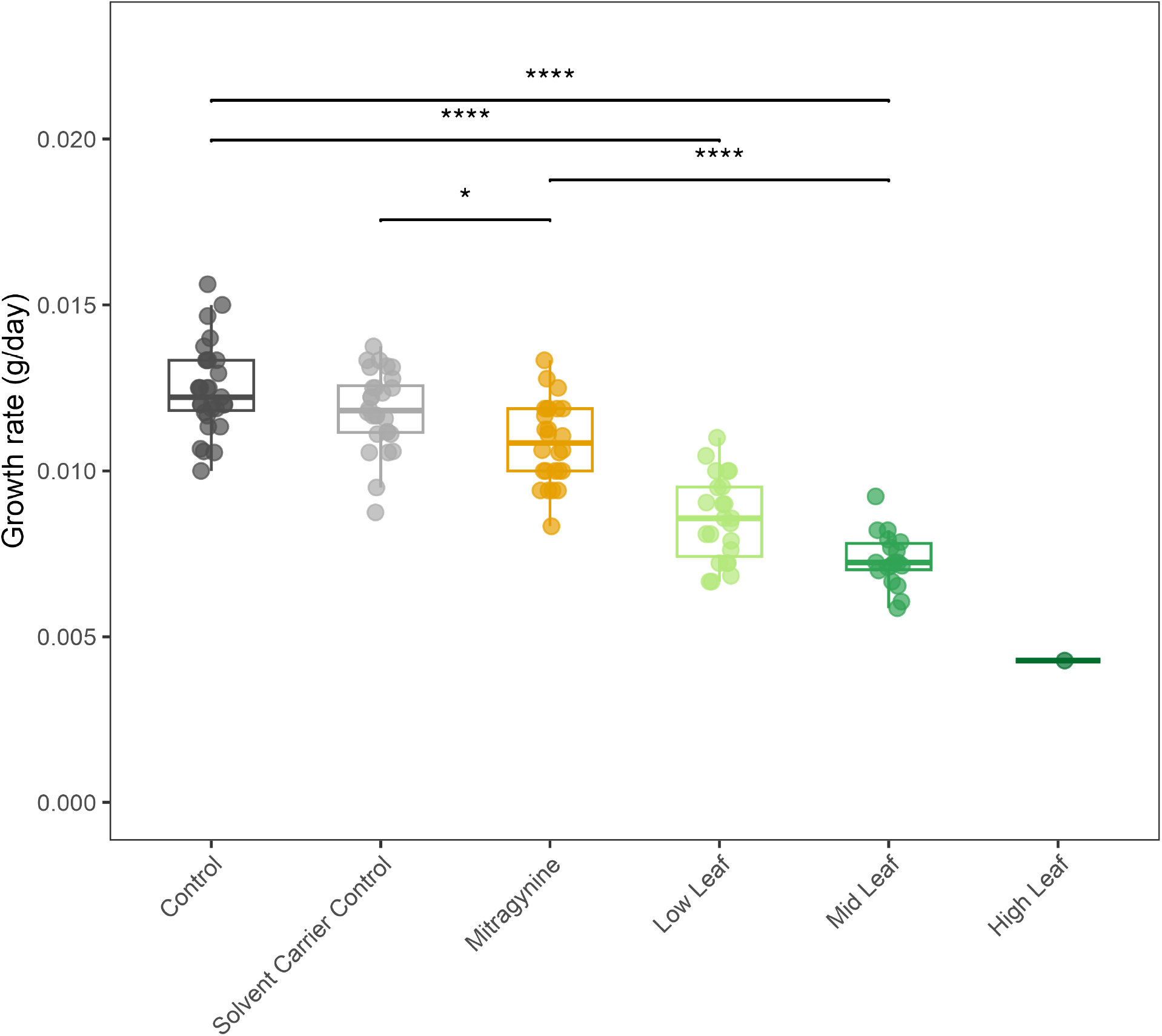
Larval growth rates. Larval growth rate (pupal weight ÷ days to pupation) by treatment group. Each point represents one individual; boxes show median and interquartile range. Brackets indicate significant pairwise differences after Holm correction (\**p* < 0.05, \*\*\*\**p* < 0.0001). Significance is not reported for comparisons to the High Leaf treatment, as only one individual survived to pupation in this group.

Larval growth rate also differed significantly across groups (Kruskal–Wallis: *H* = 87.4, df = 5, *p* < 0.0001). Median growth rates were 0.0122, 0.0118, 0.0108, 0.0086, and 0.0072 g/day for control, solvent carrier control, mitragynine, low leaf, and mid leaf, respectively. Low leaf and mid leaf larvae grew approximately 32% and 41% slower than controls, respectively (Table 2). Mitragynine-treated larvae grew approximately 10% slower than their matched solvent carrier control (*p* = 0.013), indicating a negative but modest effect of the purified alkaloid. Critically, mid leaf larvae grew significantly more slowly than mitragynine-treated larvae (*p* < 0.0001). The single surviving high leaf larva had the lowest recorded growth rate of any individual (4.3 mg/day), though this comparison did not reach significance after Holm correction given *n* = 1 (*p* = 0.103).

**Table 2.** Planned pairwise comparisons: larval growth rate.

| Comparison | W | Location shift (mg/day) | 95% CI | *p* (Holm corrected) |
| --- | --- | --- | --- | --- |
| Low Leaf vs. Control | 6.5 | -3.87 | (-4.69, -3.05) | < 1e-04 |
| Mid Leaf vs. Control | 0.0 | -4.98 | (-5.68, -4.44) | < 1e-04 |
| High Leaf vs. Control | 0.0 | -7.92 | (-11.34, -5.71) | 0.103 |
| Mid Leaf vs. Mitragynine | 1.0 | -3.51 | (-4.14, -2.8) | < 1e-04 |
| Mitragynine vs. Solvent Control | 207.0 | -1.15 | (-1.74, -0.47) | 0.013 |

\* High Leaf $n = 1$ so results should be interpreted with caution; exact CI is $\sim 92\%$ rather than $95\%$ .

Among individuals that successfully pupated, the cumulative incidence of eclosion did not differ significantly across treatment groups (Gray’s test: stat = 0.1, *p* = 1.0). Once pupated, eclosion rates were high overall: control (100%), solvent carrier control (100%), mitragynine (96.2%), low leaf (95.7%), mid leaf (100%), and high leaf (100%, though again, n=1). Of the individuals that successfully eclosed, they did so within 7-11 days of pupating across all groups. These results suggest that the main treatment effect of kratom operates primarily through larval mortality rather than pupal failure.

## Discussion

Here we demonstrate that kratom (*Mitragyna speciosa*) leaf powder causes potent, dose-dependent larval mortality and growth suppression in a model insect herbivore, offering a rare experimental test of the herbivore defense hypothesis as invoked to explain the adaptive origins of psychoactive plant compounds. While our findings are broadly consistent with the herbivore defense hypothesis, they also reveal unexpected complexity: the primary psychoactive kratom alkaloid, mitragynine, is insufficient to explain the full insecticidal activity of the leaf, and its mechanism of action in insects almost certainly differs fundamentally from its mechanism in mammals.

Larvae fed diet incorporating kratom leaf powder experienced significantly elevated mortality, delayed development, and suppressed growth rates. Incorporation of 96.3 mg/g kratom leaf powder into artificial diet caused near-complete larval mortality (97% pre-pupal death), while sublethal concentrations produced fitness costs that would be expected to substantially reduce adult reproductive potential (22; 23; 24). That these effects were observed in *S. frugiperda* — one of the most polyphagous and toxin-tolerant lepidopterans known (12) — suggests that kratom leaves present a genuinely potent defensive arsenal rather than a mild deterrent against sensitive species.

However, the most significant finding of this study is not that kratom leaf is insecticidal, but that purified mitragynine substantially underperforms relative to the whole leaf matrix. Larvae fed purified mitragynine at ∼ 0.05% w/w exhibited modestly reduced growth rates relative to the solvent carrier control, but these effects were substantially weaker than those observed in larvae fed whole leaf powder at the equivalent mitragynine concentration. Mitragynine-treated larvae did not differ significantly from the solvent carrier control in survival (log-rank *p* = 0.453), in contrast to the mid leaf treatment, which caused significantly elevated mortality relative to control (log-rank *p* = 0.002). Moreover, the low feaf diet contained approximately half the mitragynine concentration of the pure alkaloid treatment yet produced three times greater growth suppression (Table 2). This result has two important implications for the herbivore defense hypothesis in the context of psychoactive phytochemicals.

First, it suggests that mitragynine is not the primary insecticidal agent in kratom leaves, or at least it is not sufficient on its own. Kratom leaves contain at least 50 additional alkaloids (8; 25), as well as terpenoids, flavonoids, and other potentially defensive compounds (26), all of which may contribute independently or synergistically to the enhanced toxicity observed in the whole-leaf treatment. This is consistent with a growing body of evidence that phytochemical *diversity* itself is a better defense against herbivores than any single compound at equivalent concentration (27; 28). From a human pharmacology perspective, these co-occurring compounds are largely considered minor constituents, but our results suggest they may be the ecologically primary ones.

Second, and more fundamentally, the molecular target of mitragynine in mammals is almost certainly absent in insects. Mitragynine’s psychoactive and analgesic effects in vertebrates are mediated primarily through agonism at *µ*-opioid receptors (MOR) (7). While insects possess opioid-like peptides and some distantly related receptor homologs (29), these are phylogenetically and pharmacologically distinct from vertebrate MORs and are not considered functional targets for exogenous opioid ligands such as mitragynine; salvinorin A (from *Salvia divinorum*); or morphine, codeine, and thebaine (from *Papaver somniferum*) (30). Mitragynine’s psychoactivity in mammals therefore cannot be explained by the conserved receptor architecture that the herbivore defense hypothesis typically invokes to link ecological function to human drug effects (5; 31; 32).

These results invite a broader reassessment of how the herbivore defense hypothesis is applied to psychoactive compounds. The hypothesis is frequently invoked with the following logic: compound *X* is psychoactive in mammals; compound *X* is present in plant (or fungal) tissue; the plant (or fungus) is chemically defended against herbivores; therefore compound *X* evolved as a defense against herbivores and its psychoactivity in humans is an evolutionary accident. Our data illustrate why this reasoning is incomplete. Kratom leaf is genuinely insecticidal, but mitragynine — the primary psychoactive constituent of kratom — is insufficient on its own to explain that defense. The psychoactive compound and the primary defensive agent need not be the same, and conflating them produces a hypothesis that treats any demonstration of plant-level insecticidal activity as vindication of a defensive origin of a specific compound. This gap is not unique to kratom: demonstrating that *Psilocybe* extracts harm *Drosophila* (6) does not establish psychoactive psilocybin specifically as the active agent, just as our whole-leaf result does not establish that mitragynine is. We encourage this burgeoning field to adopt a more rigorous evidentiary standard so that we may move toward a more mechanistic understanding of why plants synthesize compounds that reshape animal minds.

## Acknowledgments

We are grateful to Satya Swathi Nadakuduti, Larissa Laforest, and Katherine Ransden for providing plant material and generous expertise. A special thanks to Aubrey McNutt for her forbearance during data collection and curation, and to Joshua Foley and Katelyn Fox for their assistance in data collection. This work was funded by a Student-Faculty Collaborative Grant from the ETSU Office of Undergraduate Research and Creative Activities and a Faculty Startup Award from the ETSU Office of Research and Sponsored Programs.

